# A FRET biosensor, SMART, monitors necroptosis in renal tubular epithelial cells in a cisplatin-induced kidney injury model

**DOI:** 10.1101/2022.06.18.496655

**Authors:** Shin Murai, Kanako Takakura, Kenta Sumiyama, Kenta Moriwaki, Kenta Terai, Sachiko Komazawa-Sakon, Yoshifumi Yamaguchi, Tetuo Mikami, Kimi Araki, Masaki Ohmuraya, Michiyuki Matsuda, Hiroyasu Nakano

## Abstract

Necroptosis is a regulated form of cell death involved in various pathological conditions, including ischemic reperfusion injuries, virus infections, and drug-induced tissue injuries. However, it is not fully understood when and where necroptosis occurs *in vivo*. We previously generated a Forster resonance energy transfer (FRET) biosensor, termed SMART (the sensor for MLKL activation based on FRET), which specifically monitored necroptosis in human and murine cell lines *in vitro*. Here, we generated transgenic (Tg) mice that expressed the SMART biosensor in various tissues. SMART monitored necroptosis, but not apoptosis or pyroptosis, in primary cells, including peritoneal macrophages and embryonic fibroblasts. Moreover, the FRET signal was elevated in renal tubular cells of cisplatin-treated SMART Tg mice compared to untreated SMART Tg mice. Together, SMART Tg mice may provide a valuable tool for monitoring necroptosis in different types of cells *in vitro* and *in vivo*.

## Introduction

Necroptosis is a regulated form of cell death characterized by necrotic morphology. Necroptosis is involved in various pathological conditions, including tumor necrosis factor (TNF)-mediated systemic inflammatory response syndrome (SIRS), ischemic reperfusion injury, neurodegeneration, and drug-induced tissue injuries ^1, 2^. Necroptosis is induced by death ligands (e.g., TNF, Fas ligand, and TRAIL), polyinosinic-polycytidylic acid, lipopolysaccharide (LPS), interferon (IFN)α/β, IFNγ, some virus infections, and endogenous Z-form nucleic acids ^3–5^. Among the various signaling pathways that lead to necroptosis, the TNF-mediated signaling pathway is one of the most extensively investigated ^6, 7^. In most cell types, TNF induces cell survival and inflammatory cytokine expression by activating the transcription factor, nuclear factor kappa B (NF-κB). However, under certain conditions, NF-κB activation is compromised, for example, when transforming growth factor β (TGFβ)-activated kinase 1 (TAK1) or inhibitor of apoptosis proteins (IAPs) are inhibited. Under those conditions, TNF induces apoptosis or necroptosis in a context-dependent manner ^6, 7^. Of note, under normal conditions, caspase 8 suppresses necroptosis by cleaving and inactivating receptor-interacting kinase 1 (RIPK1), an essential component of the TNF-induced necroptosis pathway ^8, 9^. When caspase 8 activity is blocked, either chemically, genetically, or by viral infection, RIPK1 recruits RIPK3 via a homotypic interaction to form a necroptosis-inducing signaling complex, termed the necrosome ^10–12^. In necrosomes, RIPK3 is activated by autophosphorylation, which leads to the formation of a higher-order amyloid-like RIPK1-RIPK3 hetero-oligomer ^13, 14^. Activated RIPK3 interacts with and phosphorylates the mixed lineage kinase domain-like pseudokinase (MLKL) ^15^. Phosphorylated MLKL forms a homo-oligomer and translocates to the cell membrane to execute necroptosis ^16–19^

The ruptured membranes of dying cells release intracellular contents, which are referred to as damage-associated molecular patterns (DAMPs). DAMPs comprise various molecules, including heat shock proteins (HSPs), high mobility group protein B1 (HMGB1), ATP, and histones. Thus, DAMPs have pleiotropic functions that become active in a context-dependent manner ^20, 21^. However, it remains unclear when and where cells undergo necroptosis and release DAMPs. We previously developed a Forster resonance energy transfer (FRET) biosensor that could specifically monitor necroptosis, which we termed the sensor for MLKL activation based on FRET (SMART) ^22^. To visualize the release of DAMPs at single-cell resolution, we modified a new technology termed Live Cell Imaging for Secretion activity (LCI-S). This technology comprises a fluorescence-conjugated, antibody-based sandwich ELISA system and total inverted reflection fluorescence microscopy ^23, 24^. By combining the SMART and LCI-S technologies, we could visualize the execution of necroptosis and the release of HMGB1 (a nuclear DAMP) at single-cell resolution ^22^.

Here, we aimed to apply the SMART biosensor to *in vivo* experiments. We generated transgenic (Tg) mice that stably expressed SMART in various tissues. SMART Tg mice grew to adulthood without any apparent abnormality and were fertile. Consistent with previous results in tumor cell lines that expressed SMART, the FRET to cyan fluorescent protein (FRET/CFP) ratio was increased in peritoneal macrophages and murine embryonic fibroblasts (MEFs) from SMART Tg mice along with progression of necroptosis, but not apoptosis or pyroptosis. These results suggested that SMART specifically monitored necroptosis in primary cells. Furthermore, in a cisplatin-induced kidney injury model, the FRET/CFP ratio of renal proximal tubule cells increased in SMART Tg mice compared to untreated SMART Tg mice. Collectively, we conclude that SMART Tg mice may be a valuable tool for monitoring necroptosis *in vivo*.

## Results

### Generation of SMART Tg mice

The SMART is an intramolecular FRET biosensor. It comprises a fragment of the kinase-like domain of MLKL, placed between the enhanced cyan fluorescent protein (ECFP) sequence and the modified yellow fluorescent protein (Ypet) sequence. The fluorescent proteins serve as the FRET donor and acceptor, respectively (Fig. 1a, *upper panel*) ^22, 25^. In the presence of stimuli that induce necroptosis, RIPK3 is phosphorylated and subsequently oligomerizes, eliciting a conformational change in SMART. This conformational change increases the FRET efficiency, and the fluorescent signal changes from ECFP to Ypet (Fig. 1a, *lower panel*).

**Figure 1.**
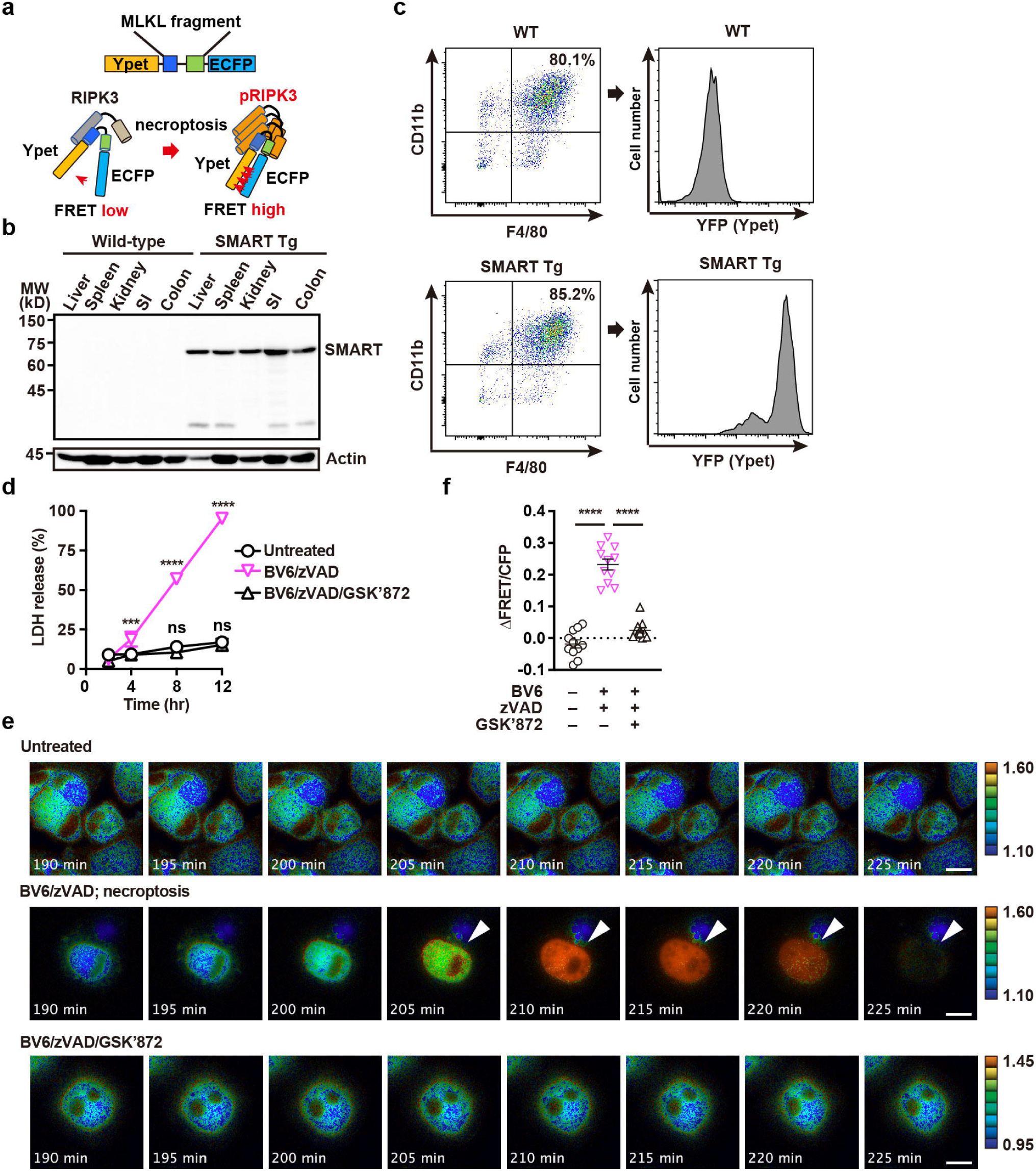
Generation of SMART Tg mice. **a**, Depiction of the SMART biosensor structure (*top*) and its activation mechanism (*bottom*). Ypet, modified yellow fluorescent protein; ECFP, enhanced cyan fluorescent protein; pRIPK3, phosphorylated RIPK3. **b,** Western blots probed with anti-GFP antibody show expression of the SMART biosensor in various murine tissues. Tissue extracts were prepared from the indicated organs of 8-week-old wild-type or SMART Tg mice. Results are representative of two independent experiments. **c**, Mice were intraperitoneally injected with thioglycollate, then peritoneal cells were recovered by washing the peritoneal cavity with ice-cold PBS on day 4 after injection. Isolated cells were stained with the indicated antibodies and analyzed with flow cytometry. The percentages of CD11b^+^F4/80^+^ cells indicate the fraction of macrophages; the levels of YFP detected in these cell populations indicate the expression of SMART. Results are representative of three independent experiments. WT, wild-type. **d**, Peritoneal macrophages from SMART Tg mice were untreated or stimulated with BV6 (1 μM) + zVAD (20 μM) or BV6 (1 μM) + zVAD (20 μM) + GSK’872 (5 μM) for the indicated times. Cell death was assessed with the LDH release assay. Results are mean ± SD of triplicate samples, and they are representative of five independent experiments. **e, f,** Peritoneal macrophages derived from SMART Tg mice were stimulated as described in **d**, and FRET/CFP ratios were calculated. Pseudocolored images show cellular changes in FRET/CFP ratio values in response to the indicated stimulations (**e**). FRET/CFP responses are color-coded according to the color scales (*right*). White arrowheads indicate cells undergoing necroptosis. Scale bars, 20 μm. Maximum changes detected in the FRET/CFP ratios (**f**). Results are mean ± SE (*n* = 11 cells per condition). Each dot indicates an individual cell. Results are representative of four independent experiments. Statistical significance was determined with two-way ANOVA with Dunnett’s multiple comparison test (**d**) or one-way ANOVA with Turkey’s multiple comparison test (**f**). \*\*\*\**P*<0.0001; ns, not significant.

To visualize necroptosis under various pathological conditions *in vivo*, we used the transposon system to generate transgenic mice that expressed SMART under the CAG promoter, as described previously ^26^. SMART Tg mice grew without any apparent abnormality, and they were fertile. These mice stably expressed SMART in various tissues, including the liver, spleen, kidney, small intestine, and colon (Fig. 1b). To test whether SMART could monitor necroptosis in primary cells, as previously demonstrated in tumor cell lines ^22^, we isolated peritoneal macrophages from SMART Tg mice. Briefly, we performed an intraperitoneal injection of thioglycollate in mice to recruit inflammatory macrophages into the peritoneal cavity; then, 4 days later, we collected the macrophages. Approximately 80% of the collected peritoneal cells expressed both CD11b and F4/80; therefore, these cells were considered macrophages (Fig. 1c, *left panel*). In addition, the expression of yellow fluorescent protein (YFP) confirmed that these macrophages expressed SMART (Fig. 1c, *right panel*).

When macrophages are treated with the IAP inhibitor, BV6, macrophages produce TNF that can result in TNF-dependent apoptosis or necroptosis in a context-dependent manner ^27, 28^. In the presence of the caspase inhibitor, zVAD, BV6 treatment induces necroptosis. When we stimulated macrophages with BV6/zVAD, the release of lactate dehydrogenase (LDH), a hallmark of membrane rupture, gradually increased. However, this increase was completely abolished when cells were treated with BV6/zVAD in the presence of the RIPK3 inhibitor, GSK’872 (Fig. 1d). Live cell imaging revealed that the FRET/CFP ratio gradually increased in cells treated with BV6/zVAD, and then it abruptly declined (Fig. 1e, f, Supplementary movie 1). The FRET/CFP ratio decline could have been due to the release of the SMART biosensor through the ruptured membrane. In contrast, the FRET/CFP did not increase in untreated cells or cells treated with BV6/zVAD/GSK’872 (Fig. 1e, f).

### SMART does not monitor pyroptosis or apoptosis

We previously reported that SMART could monitor necroptosis, but not apoptosis, in various human and murine cell lines ^22^. However, it remained unclear whether SMART could monitor pyroptosis. To address this issue, we pretreated peritoneal macrophages from SMART Tg mice with LPS, followed by nigericin stimulation. LPS plus nigericin, but not LPS alone, caused a robust release of LDH from macrophages (Fig. 2a), which suggested that the macrophages died by pyroptosis. The FRET/CFP ratio did not increase; instead, it eventually decreased in cells treated with LPS/nigericin compared to cells treated with LPS alone (Fig. 2b, c, Supplementary movie 2).

**Figure 2.**
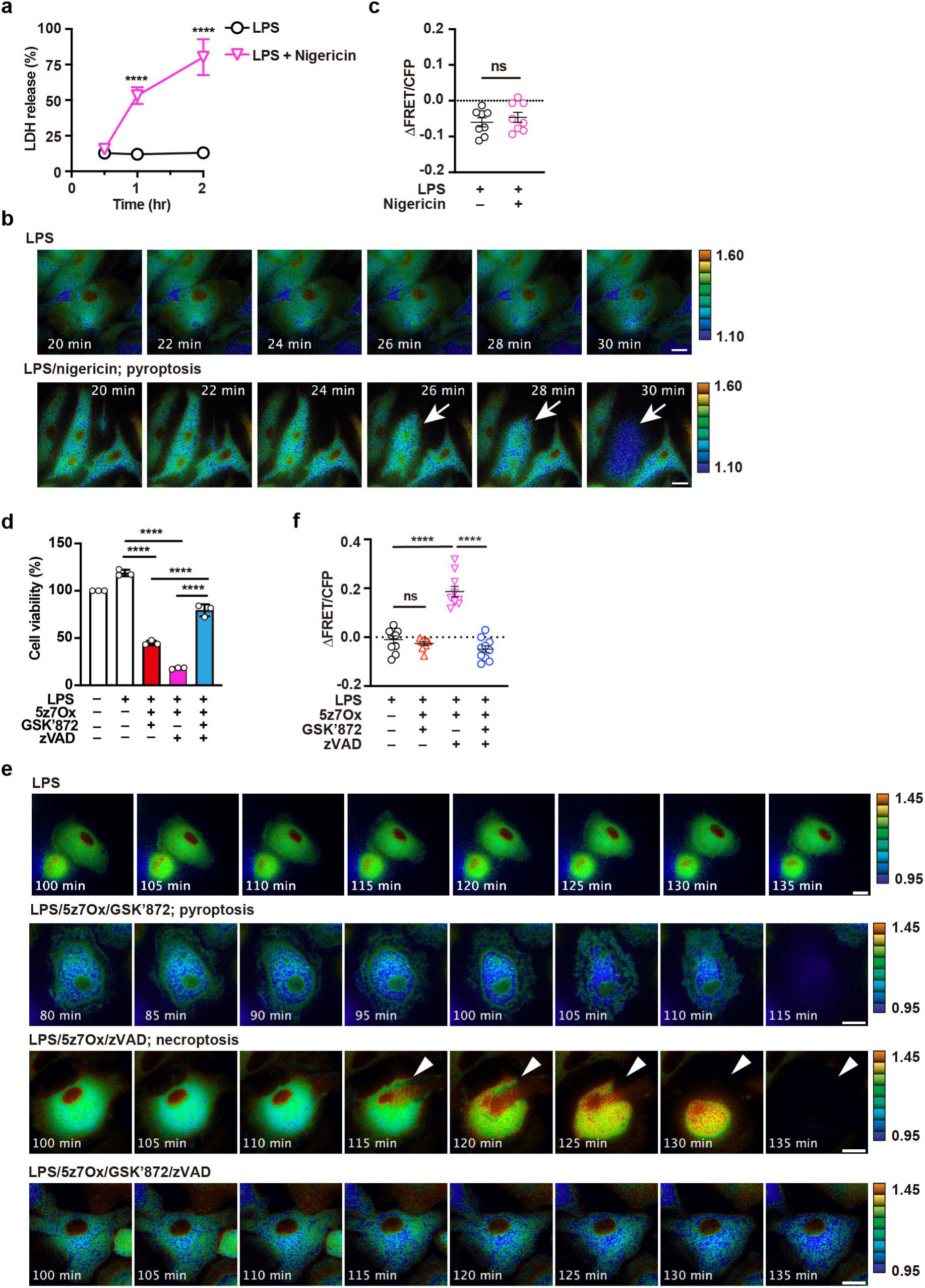
SMART does not monitor pyroptosis or apoptosis. **a**, Peritoneal macrophages isolated from SMART mice were pretreated with LPS (1 ng/mL) for 4 h, then stimulated with nigericin (10 μM) for the indicated times. Cell death was determined with the LDH release assay. Results are mean ± SD of triplicate samples and representative of three independent experiments. **b, c**, Peritoneal macrophages were stimulated as described in **a**, and the FRET/CFP ratio was calculated. Pseudocolored images show cellular changes in FRET/CFP ratio values in response to the indicated stimulations (**b**). FRET/CFP responses are color-coded according to the color scales (*right*). White arrows indicate cells undergoing pyroptosis. Scale bars, 20 μm. Maximum changes detected in the FRET/CFP ratios (**c**). Results are mean ± SE (*n* = 8 cells per condition). Each dot indicates an individual cell. Results are representative of three independent experiments. Statistical analysis was performed with the unpaired two-tailed Student’s *t*-test. ns, not significant. **d**, Peritoneal macrophages from SMART Tg mice were pretreated with LPS (1 ng/mL) for 4 h, then stimulated with the indicated combination of agents (125 nM 5z7Ox, 5 μM GSK’872, 20 μM zVAD) for 4 h. Cell viability was determined with the WST assay. Results are mean ± SD of triplicate samples. Representative results of four independent experiments. **e, f,** Peritoneal macrophages were stimulated as described in **d**. Macrophages are displayed as pseudocolor images to show cellular changes in FRET/CFP ratio values in response to the indicated stimulations (**e**). FRET/CFP responses are color-coded according to the color scales (*right*). White arrowheads indicate cells undergoing necroptosis. Scale bars, 20 μm. Maximum changes detected in the FRET/CFP ratios of the cells (**f**). Results are mean ± SE (*n* = 10 cells per condition). Each dot indicates an individual cell. Results are representative of two independent experiments. Statistical significance was determined with two-way ANOVA with Sidak’s multiple comparison test (**a**), the unpaired two-tailed Student *t*-test (**c**), or one-way ANOVA with Tukey’s multiple comparison analysis (**d, f**). \*\*\*\**P*<0.0001; ns, not significant.

Priming macrophages with LPS and subsequently stimulating them with the TAK1 inhibitor, 5z-7-oxozeaneaeol (5z7Ox), can result in apoptosis, necroptosis, or pyroptosis, depending on the context ^29–31^. To induce necroptosis selectively, we primed macrophages with LPS, then stimulated them with 5z7Ox in the presence of zVAD. In parallel experiments, after priming, we stimulated macrophages with 5z7Ox in the presence of GSK’872 to induce apoptosis or pyroptosis. Macrophage viability decreased under both of the latter conditions, but viability was maintained when cells were stimulated with LPS/5z7Ox/GSK’872/zVAD, which blocked apoptosis, necroptosis, and pyroptosis (Fig. 2d). Of note, the FRET/CFP ratio increased in cells stimulated with LPS/5z7Ox/zVAD (necroptotic condition), but not in cells treated with LPS/5z7Ox/GSK’872 (apoptotic or pyroptotic condition) (Fig. 2e, f). These results suggested that SMART could monitor necroptosis, but not apoptosis or pyroptosis, in primary macrophages.

### SMART monitors necroptosis in a RIPK3-dependent manner

Necroptosis depends on the activities of RIPK3 and MLKL ^3, 32^. To further substantiate our finding, we tested whether SMART could monitor RIPK3-dependent necroptosis in primary macrophages, as observed in tumor cell lines ^22^. To that end, we generated SMART Tg mice with a *Ripk3-/-* genetic background (SMART *Ripk3-/-* mice). We prepared peritoneal macrophages from SMART *Ripk3-/-* mice. In contrast to macrophages from SMART WT mice, BV6/zVAD-induced LDH release was abrogated in macrophages from SMART *Ripk3-/-* mice (Fig. 3a). Accordingly, the increase in the FRET/CFP ratio was abolished in macrophages from SMART *Ripk3-/-* mice after BV6/zVAD stimulation compared to those from SMART WT mice (Fig. 3b, c). These results showed that increased FRET/CFP ratio in primary macrophages depended on RIPK3.

**Figure 3.**
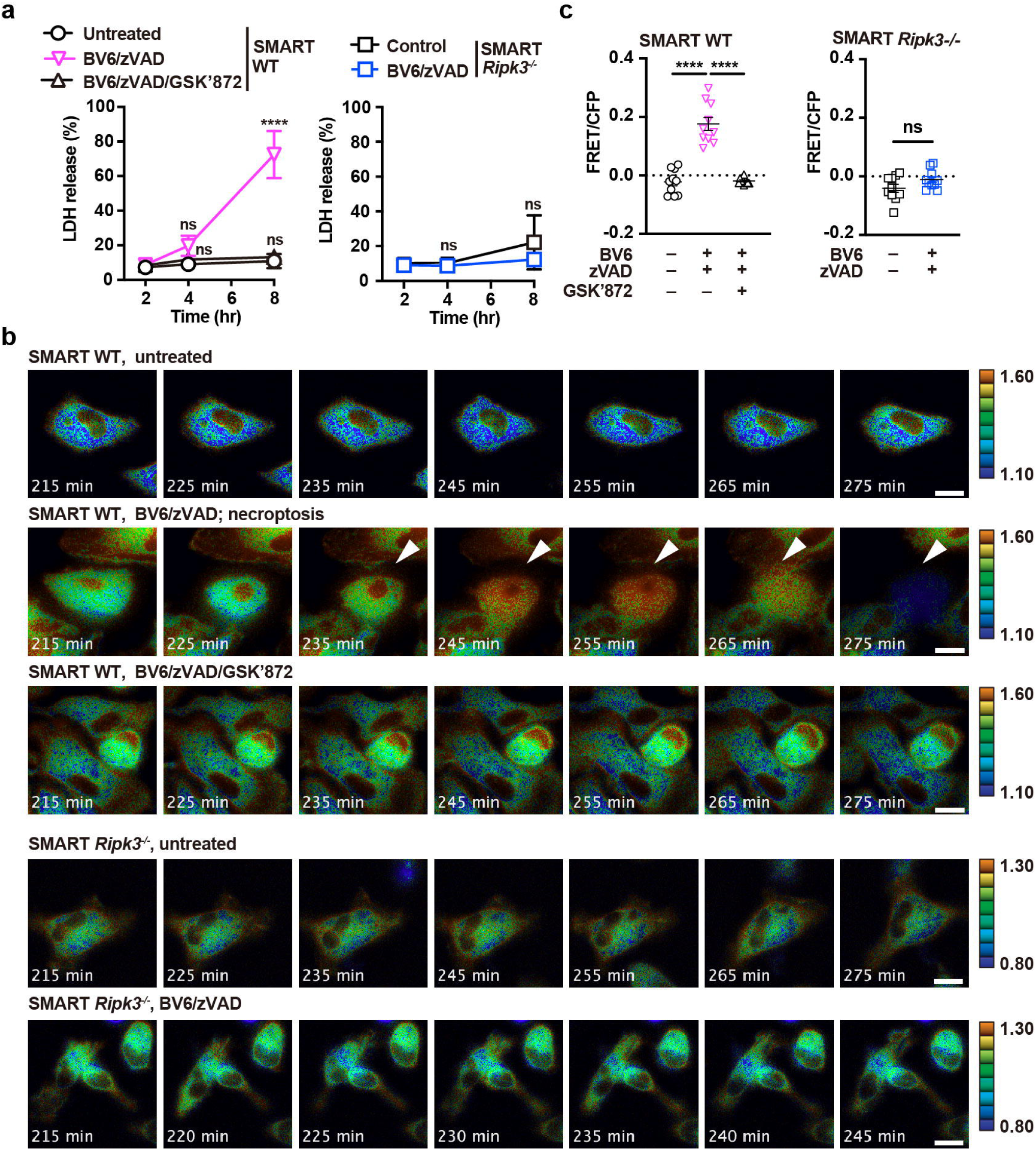
SMART monitors necroptosis in a RIPK3-dependent manner. **a**, Peritoneal macrophages from SMART WT or SMART *Ripk3^-/-^* mice were untreated or stimulated with BV6 (1 μM) + zVAD (20 μM) or BV6 (1 μM) + zVAD (20 μM) + GSK’872 (5 μM) for the indicated periods. Cell death was determined with the LDH release assay. Results are mean ± SD of triplicate samples. **b, c**, Peritoneal macrophages were stimulated and analyzed as described in **a**. Pseudocolored images show cellular changes in FRET/CFP ratio values in response to the indicated stimulations (**b**). FRET/CFP responses are color-coded according to the color scales (*right*). White arrowheads indicate cells undergoing necroptosis. Scale bars, 20 μm. Maximum changes detected in the FRET/CFP ratio (**c**). Results are mean ± SE (*n* = 10 cells). Each dot indicates an individual cell. Statistical analysis was performed with two-way ANOVA with Tukey’s multiple comparisons test (**a**, *left panel*) or Sidak’s multiple comparison test (**a**, *right panel*), one-way ANOVA with Tukey’s multiple comparison test (**c**, *left panel*), or the unpaired two-tailed Student *t*-test (**c**, *right panel*). \*\*\*\**P* <0.0001; ns, not significant. All results are representative of three independent experiments.

### SMART monitors necroptosis, but not apoptosis, in MEFs

To extend our observations in peritoneal macrophages to other cell types, we next stimulated primary MEFs derived from SMART Tg mice. We found that TNF/BV6/zVAD stimulation did not cause cell death in primary MEFs (pMEFs), but did cause cell death in immortalized MEFs (iMEFs) (Fig. 4a, b). We noticed that MLKL expression was much lower in pMEFs than in iMEFs (Fig. 4c). Because MLKL expression is induced by interferons ^33, 34^, we treated pMEFs with IFNβ. MLKL expression was strongly enhanced; accordingly, IFNβ-pretreated pMEFs died in response to TNF/BV6/zVAD stimulation (Fig. 4d). In addition, the FRET/CFP ratio was increased in IFNβ-pretreated pMEFs after TNF/BV6/zVAD stimulation, but this increase was abolished in the presence of GSK’872 (Fig. 4e, f, Supplementary movie 3). Moreover, when IFNβ-pretreated pMEFs were stimulated with TNF/BV6/GSK’872, apoptosis was induced, but the FRET/CFP ratio was not increased (Fig. 4g, h). This result suggested that SMART did not monitor apoptosis in pMEFs.

**Figure 4.**
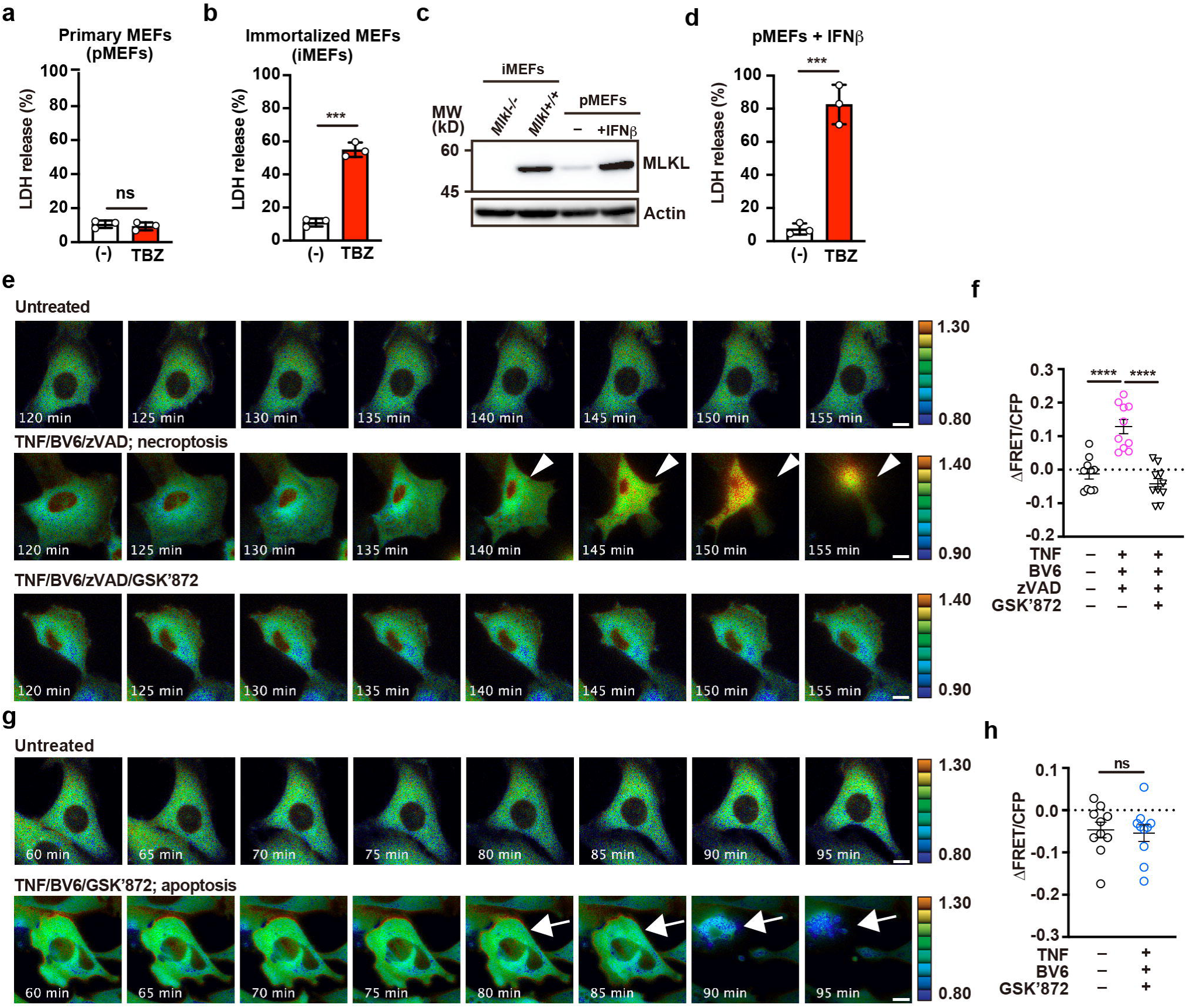
SMART monitors necroptosis, but not apoptosis, in MEFs. **a, b**, primary MEFs (pMEFs) (**a**) and immortalized MEFs (iMEFs) (**b**) were untreated or stimulated with TBZ (10 ng/mL TNF, 1 μM BV6, 20 μM zVAD) for 12 h. MEFs were generated from SMART Tg mice, and MEFs with less than five passages were used as pMEFs. Cell death was determined with the LDH release assay. Results are mean ± SD of triplicate samples. **c**, Western blots of lysates from untreated iMEFs from *Mlkl-/-* or wild-type mice, and untreated or IFNβ (2,000 IU/mL)-treated pMEFs from SMART Tg mice. Blots were probed with anti-MLKL and anti-actin antibodies. **d**, pMEFs were pretreated with IFNβ, and then stimulated with TBZ. Cell viability was determined with the LDH release assay. Results are mean ± SD of triplicate samples. **e-h**, pMEFs were untreated or stimulated with the indicated combination of agents (10 ng/mL TNF, 1 μM BV6, 20 μM zVAD, 5 μM GSK’872), and the FRET/CFP ratios were calculated. Pseudocolored images show cellular changes in the FRET/CFP ratio values in response to the indicated stimulations (**e, g**). FRET/CFP responses are color-coded according to the color scales (*right*). White arrowheads and white arrows indicate cells undergoing necroptosis (**e**) and apoptosis (**g**), respectively. Scale bars, 20 μm. Maximum changes were detected in the FRET/CFP ratios (**f, h**). Results mean ± SE (*n* = 10 cells per condition). Each dot indicates an individual cell. Statistical analyses were performed by the unpaired two-tailed Student’s *t*-test (**a, b, d, h**) or one-way ANOVA with Tukey’s multiple comparison test (**f**). \*\*\**P*<0.001; \*\*\*\**P*<0.0001; ns, not significant. All results are representative of at least two independent experiments.

### Administration of cisplatin induces necroptosis in renal proximal tubular cells

We tested whether *in vivo* imaging of necroptosis was feasible with SMART Tg mice. Previous studies have shown that necroptosis occurred *in vivo* in murine models of TNF-induced SIRS, ischemic reperfusion injuries, and cisplatin-induced kidney injuries ^35–37^. After considering which models and organs would be most suitable and technically feasible for imaging necroptosis *in vivo* with two-photon excitation microscopy, we focused on cisplatin-induced kidney injury.

Previous studies reported that administration of cisplatin induces renal proximal tubular cell necroptosis, and this injury is attenuated in *Ripk3-/-* mice ^37, 38^. We found that cisplatin injections resulted in gradual increases in blood urea nitrogen (BUN) and serum creatinine levels in wild-type (WT) mice (Fig. 5a). These signs were accompanied by kidney damage, such as the dilatation of the proximal tubule lumens (Fig. 5b). We then investigated whether phospho-RIPK3 (pRIPK3)-positive cells, a hallmark of necroptosis, appeared in the kidney after cisplatin injection. We found that, after the cisplatin injection, the numbers of pRIPK3-positive cells gradually increased and peaked on day 2 in the renal cortex of wild-type mice (Fig. 5c, e). We also found cells that cleaved caspase 3 (CC3)-positive cells appeared in the kidneys of wild-type mice after the cisplatin injection (Fig. 5d, f). These findings suggested that cisplatin-induced both apoptosis and necroptosis in the kidney. In contrast to the previous studies ^37, 38^, cisplatin-induced kidney injury was not attenuated, but rather exacerbated in *Ripk3^-/-^* mice compared to WT mice (Fig. 5g). Of note, pRIPK3^+^ cells were completely abolished in the kidneys of *Ripk3^-/-^* mice (Fig. 5h), but the numbers of CC3^+^ cells were increased in the kidneys of *Ripk3^-/-^* mice (Fig. 6i, j).

**Figure 5.**
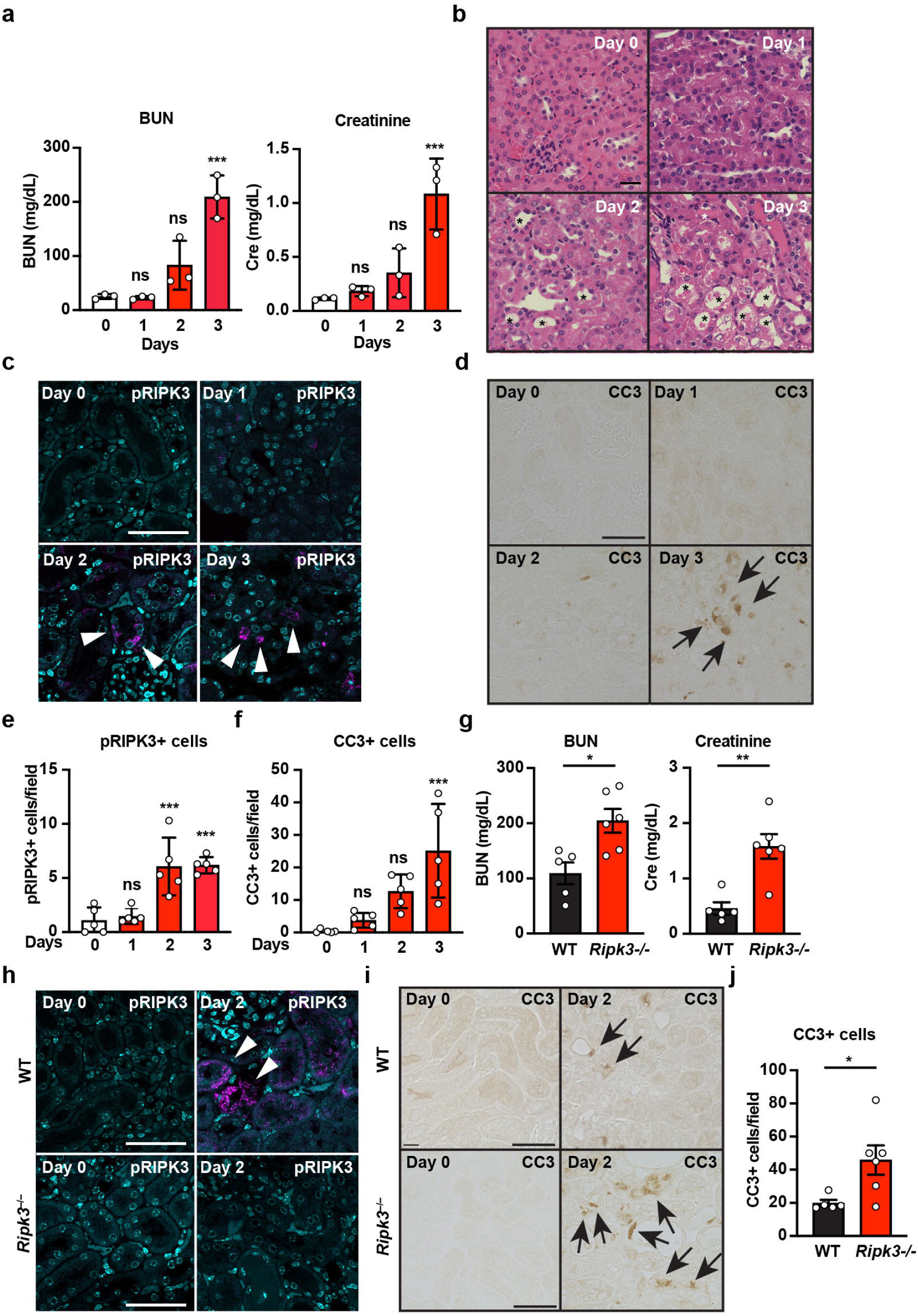
Administration of cisplatin induces necroptosis in renal proximal tubular cells. **a-f**, Eight-week-old wild-type were intravenously injected with cisplatin (20 mg/kg), then sacrificed at the indicated times. Concentrations of blood urea nitrogen (BUN) and serum creatinine were determined at the indicated times after cisplatin treatment (**a**). Results are mean ± SE (*n* = 3 mice per time point). Results are representative of two independent experiments. Kidney tissue sections were prepared at the indicated times after cisplatin treatment and stained with hematoxylin & eosin (*n* = 5 mice) (**b**). Scale bar, 50 μm. Asterisks indicate dilated proximal tubules. Kidney tissue sections were stained with anti-phospho-RIPK3 (pRIPK3) (**c**) or anti-cleaved caspase 3 (CC3) (**d**) antibodies. White arrowheads and black arrows indicate pRIPK3^+^ and CC3^+^ cells, respectively. Scale bars, 50 μm. Cell counts for pRIPK3^+^ (**e**) and CC3^+^ (**f**) cells, expressed as the number of positive cells per field. Results are mean ± SE (*n* = 5 mice). **g-j**, Eight-week-old wild-type and *Ripk3^-/-^* mice were injected with cisplatin as in **a** and sacrificed on day 2 after injection (wild-type, *n* = 5 mice; *Ripk3^-/-^* mice, *n* = 6 mice). Concentrations of BUN and serum creatinine were determined as in **a** (**g**). Kidney tissue sections were stained with anti-pRIPK3 (**h**) or anti-CC3 (**i**) antibodies. Scale bars, 50 μm. The numbers of CC3^+^ cells were counted as in **f** (**j**). White arrowheads and black arrows indicate pRIPK3^+^ and CC3^+^ cells, respectively. Statistical analyses were performed with one-way ANOVA and Dunnett’s multiple comparison test (**a, e, f**) or the unpaired two-tailed Student *t*-test (**g, j**). \*\*\**P*<0.001; ns, not significant.

**Figure 6.**
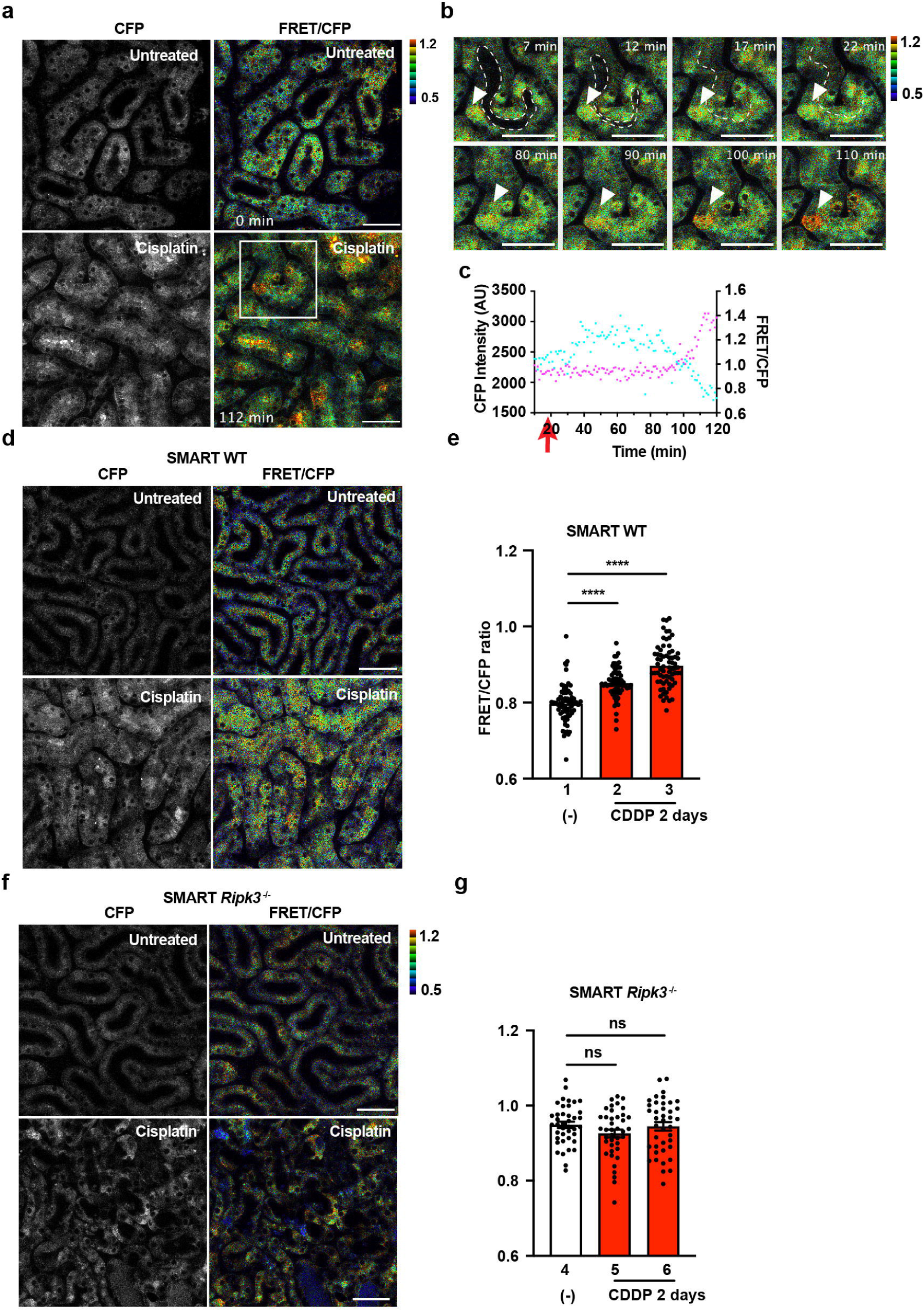
SMART monitors necroptosis in renal proximal tubular cells after cisplatin administration. **a-c**, Eight- to twelve-week-old SMART Tg mice were untreated or treated with 20 mg/kg cisplatin. On day 2, the left-side kidneys of untreated and cisplatin-treated mice were analyzed with two-photon excitation microscopy. Black-and-white and pseudocolored images show CFP intensities and the FRET/CFP ratios, respectively, in kidneys of untreated (*upper panel*) or cisplatin-treated (*lower panel*) wild-type mice (**a**). FRET/CFP responses are color-coded according to the color scales. Scale bars, 50 μm. Magnified images of the areas in **a** outlined in white boxes (**b**). Dotted white lines indicate proximal tubule lumens. White arrowheads indicate cells that show increases in the FRET/CFP ratios. FRET/CFP responses are color-coded according to the color scales. Scale bars, 50 μm. Graphs show CFP intensities (*blue*) and the FRET/CFP ratios (*pink*) of the cells indicated in **b**, calculated at the times indicated (**c**). Red arrows indicate the time that the tubule lumens were occluded. **d-g**, Eight-to twelve-week-old SMART WT mice (**d**) or SMART *Ripk3-/-* mice (**f**) were untreated or treated with cisplatin, as described in **a**. FRET/CFP responses are color-coded according to the color scales. Scale bars, 50 μm. Graphs show the averaged FRET/CFP ratios observed in tubules from untreated or cisplatin-treated SMART WT (**e**) or SMART *Ripk3-/-* (**g**) mice. Results are mean ± SE (*n* = 70 tubules in SMART WT mice; *n* = 40 tubules in SMART *Ripk3^-/-^* mice). Each number indicates an individual mouse. Results are representative of two independent experiments. Statistical analyses were performed by one-way ANOVA with Dunnett’s multiple comparison test. \*\*\*\**P*<0.0001; ns, not significant.

### SMART monitors necroptosis in renal proximal tubular cells after cisplatin administration

We then tested whether the FRET/CFP ratio was increased in SMART Tg mice with cisplatin-induced kidney injuries. Given that pRIPK3-positive cells peaked on day 2 after the cisplatin injection (Fig. 5c, e, h), we analyzed necroptosis by *in vivo* imaging on day 2. Almost all proximal tubule lumens were patent in untreated mice, and these proximal tubular cells did not show the increased FRET/CFP ratios (Fig. 6a, *upper panel*). However, in cisplatin-treated mice, many proximal tubule lumens were occluded, and several proximal tubules showed the increased FRET/CFP ratios on day 2 (Fig. 6a, *lower panel*). To investigate the kinetics of the occlusion of proximal tubule lumens and the increased FRET/CFP ratios, we visualized these two events by live imaging following cisplatin injection on day 2. In untreated mice, most proximal tubule lumens remained patent during the imaging periods (Supplementary movie 4). In sharp contrast, a few proximal tubule lumens were patent at the start of the imaging, but these tubule lumens were eventually occluded during the imaging periods (Fig. 6b, Supplementary Movie 5). The FRET/CFP ratio of proximal tubular cells was increased and peaked at approximately 100 min after the occlusion of the proximal tubule lumens (Fig. 6b, c). These results suggested that the increase in the FRET/CFP ratio may be associated with the occlusion of the tubule lumens and that SMART could monitor necroptosis of the proximal tubular cells.

To evaluate the increase in the FRET/CFP ratios in the kidneys more quantitatively, we calculated the averaged FRET/CFP ratios of each tubule in the kidneys of untreated and cisplatin-treated mice, during the same periods, on day 2 after the injection. We found significantly higher average FRET/CFP ratios in the kidneys of cisplatin-treated mice compared to the kidneys in untreated mice (Fig. 6d, e). In contrast, the averaged FRET/CFP ratios in SMART *Ripk3-/-* mice after the cisplatin injection were comparable to those observed in untreated SMART *Ripk3-/-* mice (Fig. 6f, g). Taken together, these results indicated that SMART could monitor renal proximal tubular cell necroptosis *in vivo*.

## Discussion

In the present study, we generated transgenic mice that stably expressed the FRET biosensor, SMART, in various tissues. Consistent with our previous study ^22^, we could efficiently monitor necroptosis *in vitro* in primary cells isolated from SMART Tg mice, such as macrophages and MEFs. Moreover, we could image cells in SMART Tg mice *in vivo* with two-photon excitation microscopy. We found that the FRET/CFP ratio in renal proximal tubular cells was elevated in cisplatin-treated mice compared to untreated mice. Taken together, these findings suggested that SMART Tg mice may be a valuable tool for visualizing necroptosis *in vivo*. It remains unclear whether we will be able to monitor necroptosis in all cells prepared from SMART Tg mice; nevertheless, SMART Tg mice may be useful for FRET analyses in various types of primary cells.

In contrast to apoptosis, which normally occurs in different developmental stages, necroptosis does not play a major role in development. However, necroptosis is involved in various pathological conditions, including TNF-mediated SIRS, ischemic reperfusion injuries, virus infections, neurodegeneration, and drug-induced tissue injuries ^1, 2^. Therefore, we first tested whether TNF-induced SIRS in SMART Tg mice might be a useful model for visualizing necroptosis *in vivo*. In contrast to several previous studies, which showed that intestinal epithelial cells die by necroptosis after injecting high concentrations of TNF ^35, 36^, we found that only small numbers of intestinal epithelial cells expressed pRIPK3. Moreover, small intestine peristalsis prevented live-cell imaging of necroptosis with two-photon excitation microscopy.

Considering organ accessibility from an external position, we decided to study the cisplatin-induced kidney injury model in SMART Tg mice. IHC revealed that pRIPK3^+^ cells appeared in the kidney on day 2 following cisplatin injection. This relatively delayed kinetics of the appearance of necroptotic cells prevented the standard *in vivo* imaging from determining the FRET/CFP ratio before and after cisplatin stimulation. Therefore, we started imaging the FRET/CFP ratio of proximal tubular cells in the kidneys on day 2 following cisplatin injection. Although we expected that it would be difficult to capture cells undergoing necroptosis during short-term imaging (∼ 2 h), we acquired images in SMART Tg mice that showed the increased FRET/CFP ratio in several renal proximal tubular cells. Of note, the increased FRET/CFP ratio in tubular cells usually occurred after the occlusion of proximal tubule lumens, suggesting that induction of necroptosis does not appear to be an initial event of tubular injury, but rather a late event following occlusion of proximal tubule lumens. As our present study did not investigate the causal relationship between the induction of the FRET and the occlusion of tubule lumens, we cannot formally exclude the possibility that these two events are independent, but occur simultaneously.

Of note, the numbers of cells showing the increased FRET/CFP ratios were relatively few, even in the occluded tubules of the cisplatin-treated kidneys of SMART Tg mice. This suggests that another type of cell death, such as apoptosis, ferroptosis, or necrosis, may simultaneously occur in cisplatin-induced kidney injury ^39, 40^. The mechanisms underlying cisplatin-induced necroptosis are still a matter of debate. A previous study reported that cisplatin-induced nephrotoxicity is mediated by TNF produced by renal parenchymal cells ^41^. In contrast, a later study reported that combined treatment of renal tubular cells with TNF, TNF-like weak inducer of apoptosis (TWEAK), and IFNγ, but not TNF alone, results in necroptosis ^37^. Moreover, under the conditions where expression of IAPs is attenuated by chemotherapeutic agents, such as etoposide and doxorubicin, the cell death-inducing signaling complex, termed the Ripoptosome, is spontaneously formed in the absence of death ligand stimulation ^42, 43^. The Ripoptosome comprises RIPK1, caspase 8, and FADD, thereby inducing apoptosis or necroptosis in a context-dependent manner ^42, 43^. Taken together, one might surmise that cisplatin treatment induces downregulation of the expression of IAPs that triggers the Ripoptosome formation, further increasing the susceptibility of renal proximal tubular cells to TNF- or TWEAK-induced apoptosis or necroptosis. Consistent with this idea, the RIPK1 inhibitor has been shown to suppress cisplatin-induced necroptosis ^44^. Further study will be required to investigate this possibility.

FRET/CFP ratio elevations in renal tubular cells were not observed in SMART *Ripk3-/-* mice. That finding suggested that elevations in the FRET/CFP ratios depended on RIPK3 expression. In contrast to the previous studies ^37, 44^, BUN and serum creatinine levels and numbers of CC3^+^ cells were increased in *Ripk3^-/-^* mice compared to wild-type mice following cisplatin injection, at least under our experimental conditions. Although the mechanisms underlying the discrepancy between our and previous results are currently unknown, one can surmise that DAMPs released from necroptotic tubular cells may attenuate cisplatin-induced apoptosis.

Necroptotic cells may release large amounts of intracellular content into the extracellular space, resulting in various cellular responses ^20^. Although we previously succeeded in simultaneously imaging the execution of necroptosis and the release of HMGB1 *in vitro* ^22^, it remains largely unknown where and when DAMPs are released from necroptotic cells *in vivo*. To address this issue, it will be crucial to generate transgenic mice that express DAMPs fused to a fluorescent protein, such as HMGB1-mCherry. Then, a new mouse line could be generated by crossing SMART mice with HMGB1-mCherry mice. That model might allow the visualization of DAMP release and necroptosis simultaneously, at single-cell resolution. *In vivo* imaging with SMART Tg mice could pave the way to a better understanding of necroptosis-induced biological responses *in vivo*.

## Methods

### Reagents

We purchased the following reagents as follows: BV6 (B4653, ApexBio), cisplatin (AG-CR1-3590, Adipogen), GSK’872 (530389, Merck), IFNβ (12401-1, PBL-Assay Science), LPS (434, List Labs), nigericin (AG-CN2-0020, Adipogen), Murine TNF (34-8321, eBioscience), zVAD (3188-v, Peptide Institute), and 5z7Ox (499610, Millipore). The following antibodies were used in this study: anti-actin (A2066, Sigma-Aldrich, diluted 1:1000), anti-cleaved caspase 3 (9661, Cell Signaling Technology, diluted 1:1000), phycoerythrin (PE) cyanine-7–conjugated anti-CD11b (101215, Biolegend, diluted 1:100), PE-conjugated anti-F4/80 (50-4801, TOMBO biosciences, diluted 1:100), anti-green fluorescent protein (GFP) (sc-8334, Santa Cruz Biotechnology, diluted 1:5000), anti-MLKL (3H1, Millipore, diluted 1:1000), anti-phospho-RIPK3 (57220, Cell Signaling Technology, diluted 1:1000), horseradish peroxidase (HRP)–conjugated donkey anti-rat IgG (712-035-153, Jackson ImmunoResearch Laboratories, Inc, 1:20000), HRP–conjugated donkey anti-rabbit IgG (NA934, GE Healthcare, 1:20000), biotin-conjugated donkey anti-rabbit IgG (E0432, Dako, 1:200), and Streptavidin-HRP (P0397, Dako, 1:300).

### Cell culture

Primary MEFs (pMEFs) were prepared from SMART Tg mice at E14.5 after coitus with a standard method. We prepared immortalized wild-type and *Mlkl-/-* MEFs (iMEFs) described previously ^22^. MEFs were maintained in DMEM medium containing 10% fetal bovine serum (FBS).

### Mice

*Ripk3*-/-mice^45^ (provided by Genentech Inc.) were described previously. C57BL/6 mice (Sankyo Lab Service) were housed in a specific pathogen-free facility and received a routine chow diet and water *ad libitum*. All animal experiments were performed according to the guidelines approved by the Institutional Animal Experiments Committee of Toho University School of Medicine (approval number: 21-54-400), Kyoto Graduate School of Medicine (approval number: Medkyo 21562), and the RIKEN Kobe branch (approval number: QA2013-04-11).

### Generation of SMART Tg mice

SMART Tg mice were generated by microinjecting *Tol2* mRNA and the pT2KXIG-mSMART vector into the cytoplasm of fertilized eggs from C57BL/6 mice, as described previously^26^. Eight- to twelve-week-old male and female mice were used for the *in vivo* imaging.

### Western blotting

Murine tissues were homogenized with a Polytron (Kinematica, Inc.) and lysed in RIPA buffer (50 mM Tris-HCl, pH 8.0, 150 mM NaCl, 1% Nonidet P-40, 0.5% deoxycholate, 0.1% SDS, 25 mM β-glycerophosphate, 1 mM sodium orthovanadate, 1 mM sodium fluoride, 1 mM phenylmethylsulfonyl fluoride, 1 μg/ml aprotinin, 1 μg/ml leupeptin, and 1 μg/ml pepstatin). After centrifugation, cell lysates were subjected to SDS polyacrylamide gel electrophoresis and transferred onto polyvinylidene difluoride membranes (IPVH 00010, Millipore). The membranes were immunoblotted with the indicated antibodies and developed with Super Signal West Dura Extended Duration Substrate (34076, Thermo Scientific). The signals were analyzed with an Amersham Imager 600 (GE Healthcare Life Sciences).

### Flow cytometry

Cells were stained with anti-CD11b and anti-F4/80 antibodies in flow cytometry staining buffer (eBioscience). The prepared cells were gated on forward and side scatter to identify the lymphocyte population, and then discriminating doublets. Cells were analyzed with a BD FACSCant II flow cytometer (BD Biosciences) and FlowJo software (BD Biosciences).

### Cell death assay

Macrophages were plated onto 96-well plates and cultured for 12 h in RPMI medium containing 10% FBS. Macrophages were stimulated with the IAP antagonist, BV6 (1 μM), in the presence of the apoptosis inhibitor, zVAD (20 μM), the RIPK3 inhibitor, GSK’872 (5 μM), or both, as indicated, for the times indicated. To induce pyroptosis, macrophages were pretreated with LPS (1 ng/mL) for 4 h, then stimulated with nigericin (10 μM) for the indicated times. Primary MEFs were untreated or pretreated with IFNβ (2000 IU/mL) for 24 h, then stimulated with TNF (10 ng/mL) and BV6 in the presence of zVAD (20 μM) or GSK’872 (5 μM), as indicated, for 18 h. The concentrations of LDH released from cells were determined with a Cytotoxicity Detection Kit (Roche), as described previously ^46^.

In some experiments, macrophages were pretreated with LPS (1 ng/mL) for 4 h, and then stimulated with 5z7Ox (125 nM) in the presence of zVAD (20 μM), GSK’872 (5 μM), or both, as indicated, for 4 h. Cell viability was determined with a water-soluble tetrazolium salts assay (Cell Counting kit-8, Dojindo Molecular Technologies).

### FRET analysis *in vitro*

FRET analysis was performed as previously described. Briefly, FRET signals were imaged with a DeltaVision microscope system (GE healthcare) built on an Olympus IX-71 inverted microscope base equipped with a Photometric Coolsnap HQ2 CCD camera and a 60×/NA1.516 PlanApo oil immersion lens (Olympus). For live-cell imaging FRET sensors, cells were seeded on gelatin-coated CELLview Cell Culture Dishes (Greiner Bio-One) and maintained in an incubator at 37°C with 5% CO_2_. For imaging, cells were observed with a Blue excitation filter (400-454 nm), two emission filters (blue-green, 463-487 nm for ECFP; yellow-green, 537-559 nm for Ypet), and a C-Y-m polychronic mirror. The FRET emission ratio (FRET/CFP) was calculated with SoftWoRx (Applied Precision Inc) by dividing the excitation at 436 nm and emission at 560 nm (FRET) by the excitation at 436 nm and emission at 470 nm (CFP). For statistical analyses, the obtained images were analyzed with ImageJ and MetaMorph software. The ΔFRET/CFP ratios were calculated by subtracting the FRET/CFP ratio at time 0 from the FRET/CFP ratio at the indicated times.

### Isolation of peritoneal exudate macrophages

To isolate peritoneal exudate macrophages, 6- to 8-week-old mice of the indicated genotypes were intraperitoneally injected with 2.5 ml 3% thioglycollate (T9032, Sigma). On day 4 after the thioglycollate injection, anesthetized mice were intraperitoneally injected with ice-cold PBS; then, the peritoneal cells were harvested when the PBS was recovered. This procedure was repeated twice. Harvested cells were placed in plates with RPMI medium. After removing non-adherent cells, the remaining cells were primarily peritoneal macrophages. Approximately 80% of these cells were positively stained with CD11b and F4/80 antibodies.

### Injection of cisplatin into mice

To produce cisplatin-induced kidney injuries, 8- to 12-week-old mice of the indicated genotypes were injected with cisplatin (20 mg/kg). Injected mice were sacrificed at the indicated times, and sera and kidney tissues were collected for subsequent analyses.

### Histological, immunohistochemical, and immunofluorescence analyses

Tissues were fixed in 10% formalin and embedded in paraffin blocks. Paraffin-embedded kidney sections were used for hematoxylin and eosin staining, immunohistochemistry, and immunofluorescence analyses. For immunohistochemistry, paraffin-embedded sections were treated with Instant Citrate Buffer Solution (RM-102C, LSI Medicine) for antigen retrieval. Next, tissue sections were stained with anti-CC3 antibody, followed by the secondary antibody, biotin-conjugated anti-rabbit antibody. Streptavidin-HRP was added for visualization. Images were acquired with an all-in-one microscope (BZ-X700, Keyence) and analyzed with a BZ-X Analyzer (Keyence). CC3^+^ cells were automatically counted in three randomly selected high-power fields (original magnification, ×40) per kidney, with a Hybrid Cell Count (Keyence).

For the immunofluorescence analyses, tissue sections were preincubated with MaxBlock^TM^ Autofluorescence Reducing Kit (MaxVision Biosciences), according to the manufacturer’s instructions. Next, tissue sections were stained with anti-pRIPK3 antibody, followed by visualization with the tyramide signal amplification method, according to the manufacturer’s instructions (NEL741001KT, Kiko-tech). Images were acquired with an LSM 880 (Zeiss). The images were processed and analyzed with ZEN software (Zeiss) and an image-processing package, Fiji (https://fiji.sc/). pRIPK3^+^ cells were counted manually.

### Measurement of blood urea nitrogen and creatinine

After the cisplatin injection, serum samples were collected on days 0, 1, 2, and 3. Serum creatinine (measured with an enzymatic method) and blood urea nitrogen (measured with the urease-glutamate dehydrogenase method) were measured by Oriental Yeast.

### Imaging cisplatin-induced kidney injury

Living mice were observed with an FV1200MPE-BX61WI upright microscope (Olympus) equipped with an XLPLN 25XW-MP 25X/1.05 water-immersion objective lens (Olympus). The microscope was equipped with an InSight DeepSee Ultrafast laser (0.95 Watt, 900 nm; Spectra-Physics, Mountain View, CA). The scan speed was set at 2 µs/pixel. The excitation wavelength for CFP was 840 nm. Fluorescence images were acquired with the following filters and mirrors: an infrared-cut filter (BA685RIF-3); two dichroic mirrors (DM505 and DM570); and three emission filters, including: an FF01-425/30 (Semrock, Rochester, NY), for the second harmonic generation; a BA460-500 (Olympus) for CFP; and a BA520-560 (Olympus) for FRET. The microscope was also equipped with a two-channel GaAsP detector unit and two multi-alkali detectors. FluoView software (Olympus) was used to control the microscope and acquire images. Acquired images were saved in the multilayer 12-bit tagged image file format and processed and analyzed with Metamorph software, as described previously ^47^.

Intravital mouse imaging was performed essentially as described previously ^47^. Briefly, two days before imaging, cisplatin was injected intravenously. To observe the kidney, the mouse was placed in the prone position on an electric heat pad maintained at 37℃. A 1-cm incision was made in the skin of the lower back and underlying peritoneum to expose approximately 0.25 cm^2^ of tissue. The exposed tissue was imaged with an aspiration fixation system. The obtained images were analyzed by Metamorph. The FRET/CFP ratios of relevant areas were calculated with ImageJ software. Then, the averaged FRET/CFP ratios were calculated for each proximal tubule.

### Statistical analysis

Statistical analyses were performed with the unpaired two-tailed Student’s *t*-test or one-way or two-way ANOVA with Dunnett’s, Sidak’s, or Tukey’s multiple comparison test, as appropriate. *P*-values <0.05 were considered statistically significant.

## Acknowledgments

We thank K. Asano for technical advice, Genentech Inc. for *Ripk3-/-* mice, M. Pasparakis for *Mlkl-/-* MEFs, and K. Kawakami for the transposon-based expression vector system. This work was supported, in part, by Grants-in-Aid for Scientific Research (B) 20H03475 (to HN), Scientific Research (C) 19K07399 (to KM), and Scientific Research (C) 20K05238 (to SM) from the Japan Society for the Promotion of Science (JSPS), a Grant-in-Aid for Scientific Research on Innovative Areas (JP16H06280) ― Platforms for Advanced Technologies and Research Resources “Advanced Bioimaging Support”, from the Japan Agency for Medical Research and Development (AMED), under Grant Numbers 21gm1210002 (to HN) and 21wm0325050 (to KM), grants from the Ministry of Education, Culture, Sports, Science, and Technology, Japan, Toho University Grant for Research Initiative Program (TUGRIP) (to HN), the Science Research Promotion Fund, and The Promotion and Mutual Aid Corporation for Private Schools of Japan (to HN), a GSK Japan Research Grant 2020 (to KM), and a grant from the Takeda Science Foundation (to KM).

## Author Contributions

S.M., K.M., Kenta T., M.M., and H.N. designed research; S.M., Kanako T., and S.K.-S. performed research; K.S., K.A., and M.O. contributed to new reagents/analytical tools; S.M., Kanako T., K.M., Kenta T., Y.Y., T.M., M. M., and H.N. analyzed data; S.M. and H.N. wrote the manuscript.

## Biological materials availability

All the biological materials, including SMART Tg mice used in this study, are available from the corresponding authors upon reasonable request. Obtaining *Ripk3-/-* mice requires Material Transfer Agreement (MTA) from Genentech Inc.

## Data availability

The authors declare that the data supporting this study are available within the paper and its supplementary movies. Source data behind the graphs are available as source data files. Other datasets generated during and/or analyzed during the current study are available from the corresponding author upon reasonable request.

