## Supplementary Information for "A FRET biosensor, SMART, monitors necroptosis in renal tubular epithelial cells in a cisplatin-induced kidney injury model"

**Supplementary Movies 1-5**

**Supplementary Movie 1.** Imaging of necroptosis in peritoneal macrophages, Related to Figure 1

Peritoneal macrophages from SMART Tg mice were stimulated with BV6/zVAD.

**Supplementary Movie 2.** Imaging of pyroptosis in peritoneal macrophages

Related to Figure 2

Peritoneal macrophages from SMART Tg mice were primed with LPS, then stimulated with nigericin.

**Supplementary Movie 3.** Imaging of necroptosis in MEFs, Related to Figure 4

MEFs from SMART Tg mice were stimulated with TNF/BV6/zVAD.

**Supplementary Movie 4.** Imaging of the renal proximal tubular cells in the kidney of untreated SMART Tg mice, Related to Figure 6

SMART Tg mice were untreated, and the kidney was analyzed by two-photon excitation microscopy. The time (min) since the movie started on day 2 is shown.

**Supplementary Movie 5.** Imaging of the renal proximal tubular cells in the kidney of CDDP-treated SMART Tg mice, Related to Figure 6

SMART Tg mice were treated with CDDP, and then the kidney was analyzed by two-photon excitation microscopy on day 2 following CDDP injection. The time (min) since the movie started on day 2 is shown.
